# Face processing of social cognition in patients with first episode psychosis: Its deficits and association with the right subcallosal anterior cingulate cortex

**DOI:** 10.1101/2021.03.12.435039

**Authors:** Zui Narita, Hironori Kuga, Peeraya Piancharoen, Andreia Faria, Marina Mihaljevic, Luisa Longo, Semra Etyemez, Ho Namkung, Jennifer Coughlin, Gerald Nestadt, Frederik Nucifora, Thomas Sedlak, Rebecca Schaub, Jeff Crawford, David Schretlen, Koko Ishizuka, Jun Miyata, Kun Yang, Akira Sawa

## Abstract

The clinical importance of social cognition is well acknowledged in patients with psychosis, in particular those with first episode psychosis (FEP). Nevertheless, its brain substrates and circuitries remain elusive, lacking precise analysis between multimodal brain characteristics and behavioral sub-dimensions within social cognition. In the present study, we examined face processing of social cognition in 79 FEP patients and 80 healthy controls (HCs). We looked for a possible correlation between face processing and multimodal MRI characteristics such as resting-state functional connectivity (rsFC) and brain volume. We observed worse recognition accuracy, longer recognition response time, and longer memory response time in FEP patients when compared with HCs. Of these, memory response time was selectively correlated with specific rsFCs, which included the right subcallosal sub-region of BA24 in the ACC (scACC), only in FEP patients. The volume of this region was also correlated with memory response time in FEP patients. The scACC is functionally and structurally important in FEP-associated abnormalities of face processing measures in social cognition.

## Introduction

Social cognition is the ability to use external information in social contexts to regulate interpersonal functioning, resulting in effective social behavior. Social cognition has become a high priority area in the field of psychiatry as evidenced by a growing empirical literature and increased interests in the research community (Couture et al., 2006; Green et al., 2019, 2015, 2008, 2005). Importantly, social cognition is reportedly predictive of quality of life and functional outcomes of patients with schizophrenia (Harvey and Penn, 2010; Henry et al., 2016). In a NIMH workshop, five dimensions in social cognition were underscored: *theory of mind, social perception, social knowledge, attributional bias*, and *emotional processing* (Green et al., 2008). Distinct batteries are available for each dimension in social cognition (Couture et al., 2006; Green et al., 2008, 2005). Face processing task is extensively studied for the *emotional processing* dimension, and is reportedly a promising indicator for some mental disorders, when compared with healthy controls (Gur et al., 2017).

Face processing task is performed by showing multiple faces to subjects in order to commonly evaluate accuracy and response time (Carter et al., 2009; Gur et al., 2017, 2006; Irani et al., 2012; Pinkham et al., 2018). The Karolinska Directed Emotional Faces (KDEF) is a set of human faces frequently used in research (Calvo and Lundqvist, 2008; Goeleven et al., 2008; Lundqvist et al., 1998). In addition to the KDEF, several other face processing tasks are available. For example, the Pictures of Facial Affect is the first face processing task that is widely recognized (Ekman and Friesen, 1976; Holt et al., 2005; Rasetti et al., 2009). The Penn Computerized Neurocognitive Battery and the Penn Emotion Recognition Task have also been used, particularly in psychosis research (Butler et al., 2009; Carter et al., 2009; Gur et al., 2017; Irani et al., 2012; Pinkham et al., 2018).

Several studies have pointed out that patients with schizophrenia showed impaired capacity in face processing, compared with healthy controls (HCs) (Corcoran et al., 2015; Gur et al., 2017, 2006; Kohler et al., 2010). The amygdala is one of the most intensively studied regions in task-based functional MRI (fMRI) in association with face processing, but findings remain inconclusive, with decreased (Barbour et al., 2010; Habel et al., 2004), increased (van Buuren et al., 2011), or unchanged (Cao et al., 2016; Rasetti et al., 2009) activity/connectivity in patients compared with HCs. Recently, a few resting-state fMRI (rs-fMRI) studies evaluated patients with schizophrenia along with face processing tasks (Maher et al., 2019; Martínez et al., 2019). In these studies, candidate-region approaches were employed, showing a correlation between face processing measures and resting-state functional connectivity (rsFC) of pulvinar and fusiform face areas (Maher et al., 2019; Martínez et al., 2019). Nevertheless, to the best of our knowledge, no study has addressed possible correlation between rsFCs and face processing measures in an unbiased approach for patients with psychosis.

In the present study, we studied 79 patients with first episode psychosis (FEP) and 80 HCs to evaluate possible deficits of face processing and associated brain abnormalities through a multimodal MRI assessment that includes rs-fMRI and structural MRI. We first assessed whether the FEP group was different from HCs in face processing as measured by the KDEF. We then addressed possible abnormalities in the cerebral cortex and deep grey matter associated with deficits in face processing by examining rs-fMRI data in an unbiased manner to pin down promising brain region candidates, which was followed by an analysis of structural MRI data for the specific brain region.

## Methods

### Participants

This study was approved by the Johns Hopkins School of Medicine Institutional Review Board and conducted in accordance with the Code of Ethics of the World Medical Association. Written informed consent was obtained from all adult participants aged 15 years or older. Parental consent and assent were obtained for subjects younger than 18 years. Patients with the onset of psychosis within 24 months of the study and HCs were recruited by the Johns Hopkins Schizophrenia Center. We call them as FEP patients in the present study to differentiate our cohort from recent onset psychosis cohorts in which patients with a longer duration of illness (e.g., onset within 5 years) are usually included (Deakin et al., 2018; Koshiyama et al., 2018). We acknowledge that a 2-year-window is a relatively relaxed definition for FEP. Nevertheless, such definition has been used in many published studies and meta-data analyses (Galderisi et al., 2009; Kim et al., 2020; Lesh et al., 2021; Wilson et al., 2020). Further details about the recruitment and eligibility criteria can be found in previous papers (Faria et al., 2019; Kamath et al., 2019, 2018; Wang et al., 2019).

The FEP patient group consists of patients with schizophrenia (*n* = 41), schizoaffective disorder (*n* = 9), bipolar disorder with psychotic features (*n* = 18), major depressive disorder with psychotic features (*n* = 6), psychotic disorder: not otherwise specified (NOS) (*n* =2), and schizophreniform disorder (*n* = 3).

### Face processing tasks

The KDEF is a validated image set with 4,900 standardized pictures of human faces with different emotional expressions portrayed by 70 different White individuals (Calvo and Lundqvist, 2008; Goeleven et al., 2008; Lundqvist et al., 1998). Twenty-four faces from six models (three females and three males) were randomly selected out of the KDEF set, and were used to evaluate *emotional processing*.

Face processing tasks contained one of four expressions: happy, angry, sad, or neutral. Faces were presented one at a time against a gray background in the center of the computer. Each trial started with the presentation of a face, and the face stayed in view for a maximum time of 4.5 seconds. Two subtasks were conducted: facial affect recognition task and face memory task. For the facial affect recognition task, participants were instructed to identify the expression of each face as accurately and quickly as possible to measure the response time in a valid manner. In the present study, participants were asked to press one of four labeled keys that corresponded to one of the four expressions. Each face was shown only once during the task and the maximum duration of this task was 10 mins. The face memory task was then performed, which displayed 24 faces that had already been presented and 24 new faces to participants (24 females and 24 males). Participants were asked to select whether each face had appeared in the previous task and to respond as accurately and quickly as possible by pressing one of two labeled keys (keys for yes or no). There was no time limit for this task. The interval between the facial affect recognition task and the facial memory task was five minutes. We collected the data from four outcome measures: (i) recognition accuracy, (ii) recognition response time, (iii) memory accuracy, and (iv) memory response time.

Each task was performed out of the MRI scanner. For both subtasks, greater accuracy score and shorter response time suggest better *emotional processing*.

### MRI imaging

We acquired the synchrony of fMRI time courses and the T1-WI images from cortex and deep grey matter (average volume: 1254.2 ml). The average duration between the MRI scan and social cognitive tasks was 11.2 days. The image parameters were (i) rs-fMRI: axial orientation, original matrix 80×80, 36 slices, voxel size of 3×3×4 mm, TR/TE 2000/30 ms, and 210 dynamics and (ii) T1 high-resolution-weighted images (T1-WI): sagittal orientation, original matrix 170×170, 256 slices, voxel size 1×1×1.2 mm, TR/TE 6700/3.1 ms. We chose to examine the rs-fMRI signal between pairs of regions of interest (ROIs) for several reasons. This method reduced the dimensions of the data (from voxels to ROIs) by employing “anatomical” filters. These filters were the structures in question, whose definitions were based on previous biological knowledge. Therefore, the interpretation of the results, and possible clinical translation was more straightforward (Faria et al., 2017). This method also facilitated the combination of multiple features (Faria et al., 2012) such as fMRI correlations and other variables. This was ideal in a multidimensional approach for biological research. This method was also reliable and robust against artifactual noise (Liang et al., 2015).

The segmentation involved orientation and homogeneity correction in MRICloud (www.MRICloud.org); two-level brain segmentation (skull stripping, then whole brain); image mapping based on a sequence of linear, non-linear algorithms, and Large Deformation Diffeomorphic Mapping; and a final step of multi-atlas labeling fusion, adjusted by PICSL (Tang et al., 2013). We used the multi-atlas set “Adult22_50yrs_283Labels_26atlases_M2_252_V9B.” rs-fMRI data were also processed in MRICloud (Faria et al., 2012). This processing consisted of co-registering the T1-WI and the respective segmentations to the motion and slice timing-corrected resting-state dynamics. Time courses were extracted from all cortical and subcortical gray matter regions, as defined in the atlases and regressed for physiological nuisance. The MRICloud pipeline includes well accepted protocols to minimize artifacts in the images and in the nuisance correction routine. These and other technical procedures are detailed in the original MRICloud publication (Faria et al., 2012). Seed-by-seed correlation matrices were obtained from the nuisance-corrected time courses and z-transformed by the Fisher’s method. 3,003 pairwise resting-state z-correlations from 78 regions (cortical and subcortical gray matter ROIs) were used for the analyses.

Realignment and head motion evaluation were preprocessed with ART (SPM toolbox). To examine head motion, we used framewise displacement (FD) that was calculated by six motion parameters (Power et al., 2012). FD ≥ 0.3 was used as the cutoff to identify outliers (Achterberg and van der Meulen, 2019; DeSerisy et al., 2020; Li et al., 2017). Accordingly, one subject was identified as an outlier and excluded from data analysis.

### Statistical analyses

Statistical analyses were conducted using R version 3.5.3. A student t-test and a chi-square test were used to evaluate differences in demographic features between FEP patients and HCs.

General linear regression was conducted to compare face processing measures between groups with age, sex, race, smoking status (yes/no), and years of education as covariates. When we specifically assessed the effects of race on face processing measuers, among these covariates, we did not include race as one of them. General linear regression was also conducted to compare rsFCs between groups with age, sex, race, smoking status (yes/no), and handedness as covariates. The Benjamini-Hochberg procedure, a popular method for controlling the false discovery rate, was used for multiple comparison correction. P values corrected with the Benjamini-Hochberg procedure are presented as q values.

General linear regression was used to examine the correlation between face processing measures and rsFCs: age, sex, race, smoking status, years of education, and handedness were included as covariates in the analysis for HCs, while age, sex, race, smoking status, years of education, handedness, chlorpromazine (CPZ) equivalent dose estimated using the Defined Daily Doses method (Leucht et al., 2016), and duration of illness (DOI) were included as covariates in the analysis for FEP patients. The Benjamini-Hochberg procedure was used for multiple comparison correction. We didn’t include the usage of cannabis and FD as covariates directly in the linear regression analyses to avoid over-fitting and artificial effects for the following reasons: 1) the head motion has already been adjusted in the initial steps of brain imaging data processing; 2) we only have cannabis usage records from self-report, where the dose and frequency of cannabis usage may not be accurate., However, to access whether there are any effects of head motion and cannabis on rsFCs, we performed linear regression analyses with rsFC as a dependent variable, FD or cannabis usage (yes/no) as an independent variable, as well as age, sex, race, handedness, and diagnosis (patient or control) as covariates.

General linear regression was further conducted to examine the correlation between face processing measures and the volume of regions that were pinned down by the analysis of rsFCs. The same variables considered as covariates in the correlation analysis between face processing measures and rsFCs were also utilized as covariates in this analysis.

## Results

### Demographic features

The demographic features of participants were summarized in **Table 1**. The FEP group did not differ from the HC group in race and handedness, but showed significantly younger age, higher proportion of males, higher number of smokers, higher number of cannabis users, and shorter years of education. The mismatches of these demographic factors were considered and adjusted in data analyses.

**Table 1.** Demographic summary of study participants. Abbreviations: SD, standard deviation; FEP, first episode psychosis; HC, healthy control; FD, framewise displacement; CPZ, chlorpromazine; and DOI, duration of illness.

| Characteristic | Mean ± SD | | P value |
| --- | --- | --- | --- |
|  | FEP (n=79) | HC (n=80) |  |
| Age (year) | 22.53 ± 4.62 | 23.63 ± 4.01 | 0.11 |
| Sex |  |  |  |
| Male | 55 | 32 | 2.33E-04 |
| Female | 24 | 48 |  |
| Race |  |  |  |
| African American | 36 | 53 | 0.08 |
| Caucasian | 31 | 21 |  |
| Other | 12 | 6 |  |
| Smoking status |  |  |  |
| Yes | 22 | 2 | 4.06E-06 |
| No | 57 | 78 |  |
| Cannabis usage status |  |  |  |
| Yes | 21 | 7 | 3.53E-03 |
| No | 58 | 73 |  |
| Years of education | 13.19 ± 2.56 | 14.69 ± 2.08 | 8.20E-5 |
| Handedness |  |  |  |
| Right | 70 | 71 | 1.00 |
| Left | 9 | 9 |  |
| FD | 0.08 ± 0.05 | 0.07 ± 0.04 | 0.01 |
| CPZ dose (mg) | 314.22 ± 252.72 | *N/A* | *N/A* |
| DOI (month) | 14.06 ± 8.95 | *N/A* | *N/A* |

### Difference in face processing between groups

First, we tested whether the FEP group showed deficits in face processing in comparison with the HC group using general linear regression. We observed significantly worse performances in recognition accuracy (q-value = 0.03), recognition response time (q-value = 7.52E-05), and memory response time (q-value = 4.52E-05), but not in memory accuracy (q-value = 0.09) (**Fig. 1**).

**Figure 1.**
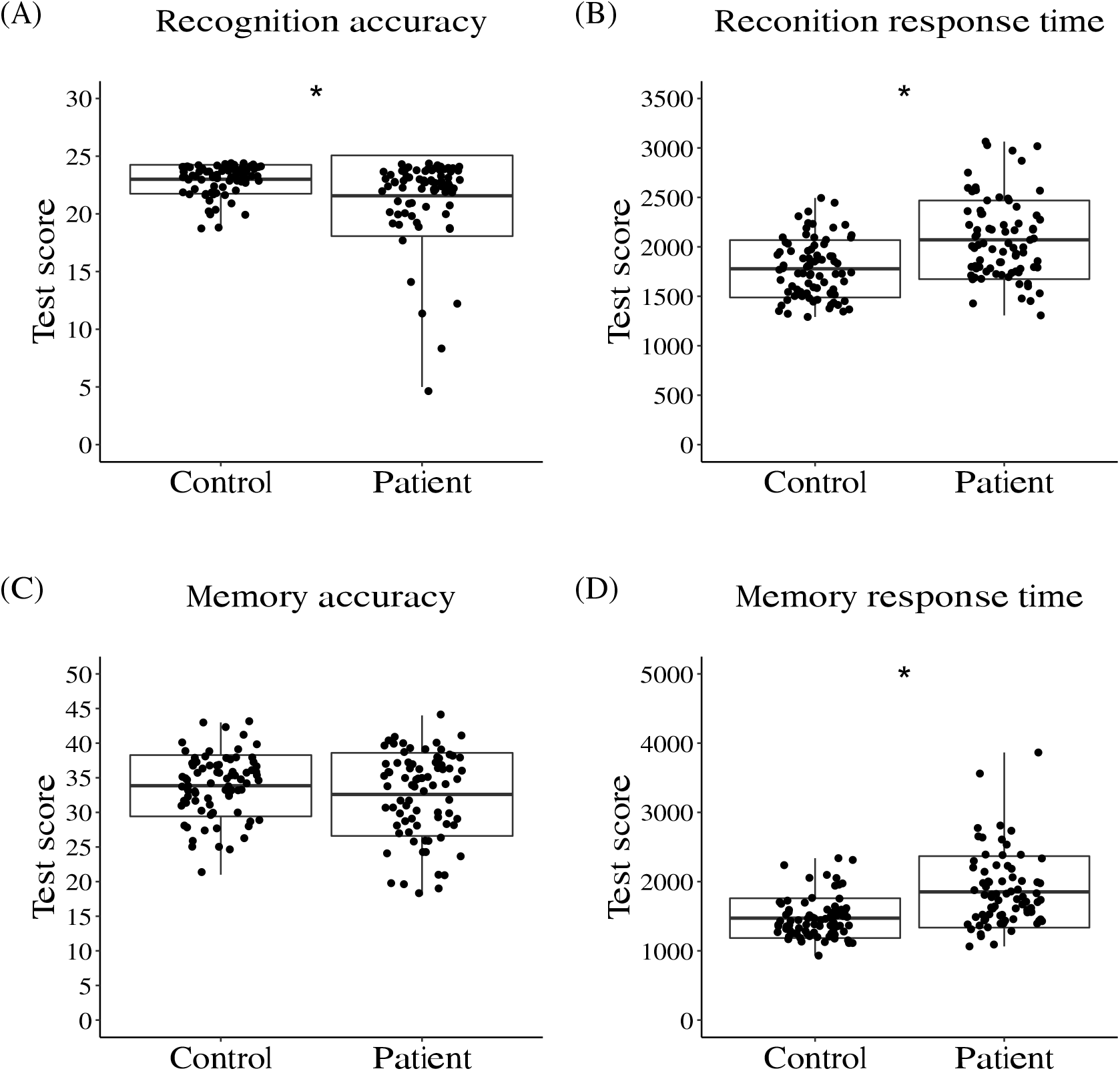
Face processing measures were significantly different between first episode psychosis (FEP) patients and healthy controls (HCs). (A) Recognition accuracy (the number of correct answers). FEP patients showed worse scores compared with HCs. (B) Recognition response time. FEP patients showed worse scores compared with HCs. (C) Memory accuracy (the number of correct answers). The difference between FEP patients and HCs didn’t reach the significance cutoff. (D) Memory response time. FEP patients showed worse scores compared with HCs. *Significant after the adjustment of age, sex, race, smoking, and years of education and multiple comparison correction (q-value < 0.05).

We questioned whether race might influence the results of face processing measures. Thus, we conducted the group comparsions between African American FEP patients and non-African American FEP patients, as well as African American HCs and non-African American HCs. We didn’t observe significant differences in any face processing measure in these group comparisons (**Supplementary Table S1**). Together, FEP patients showed deficits in KDEF-based face processing assessment.

### Possible involvement of the right subcallosal sub-region of BA24 in the anterior cingulate cortex (scACC) in memory response time in FEP patients

Then, we conducted an unbiased analysis to investigate the relationship between rsFCs and face processing. Among combinations of 3,003 rsFCs and 4 face processing measures, only the rsFC between right scACC and right precuneus was significantly correlated with memory response time after multiple comparison correction in FEP patients (q-value = 0.03) (**Fig. 2A, Supplementary Table S2**). No significant result was observed in HCs (**Supplementary Table S3**). Consistent with this finding, we also found that the rsFC between right scACC and right precuneus was significantly increased in FEP patients, compared to HCs (p-value = 0.01), in the group comparison (**Fig. 2B**).

**Figure 2.**
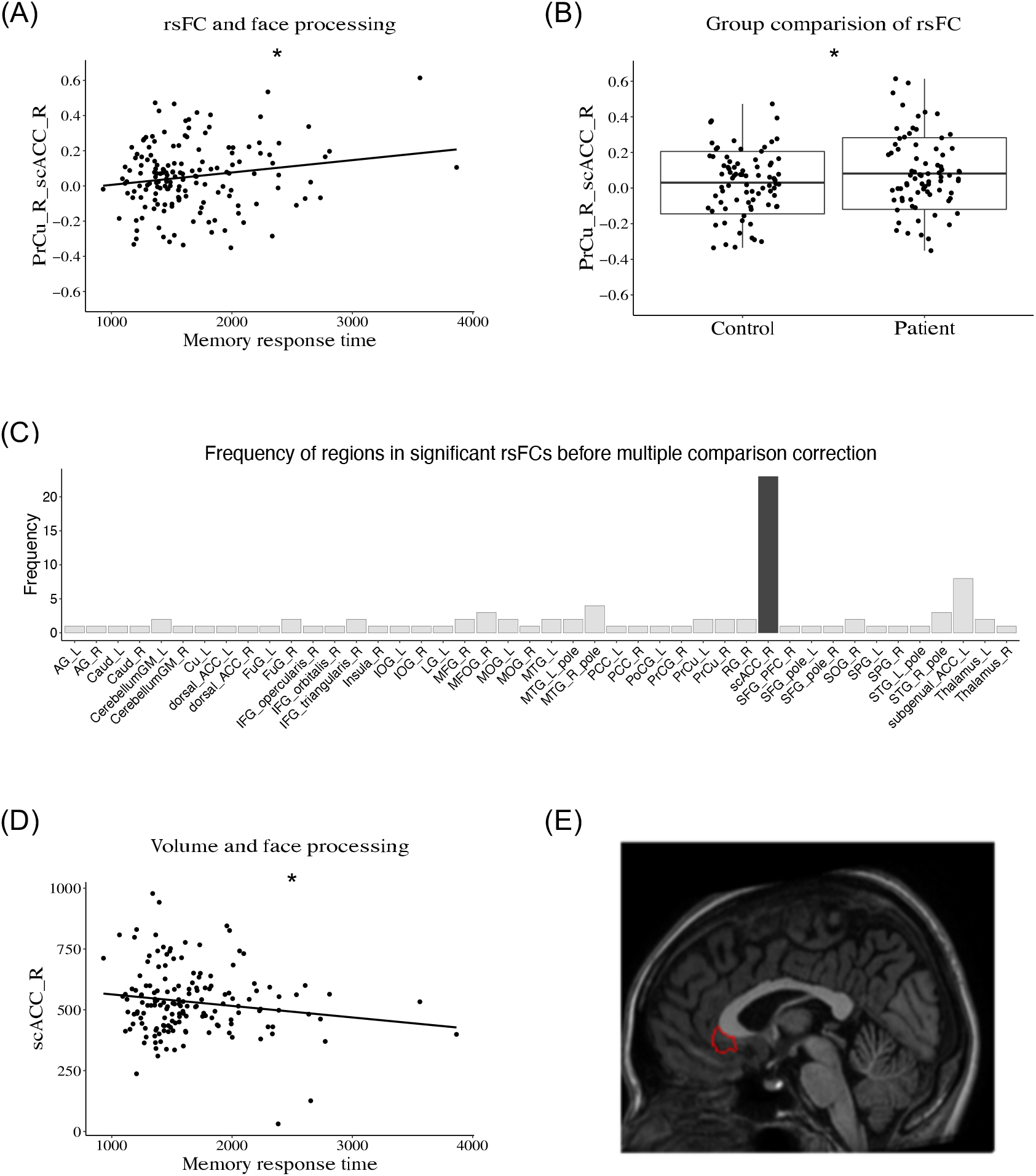
Right subcallosal sub-region of BA24 in the anterior cingulate cortex (scACC) and face processing. (A) Correlation between memory response time and resting-state functional connectivity (rsFC) between right scACC and right precuneus in FEP patients. (B) Boxplot of rsFCs between right scACC and right precuneus in FEP patients and HCs. (C) Brain regions appeared in 46 rsFCs correlated with memory response time before multiple comparison correction: scACC appeared prominently (23 rsFCs out of 46). The full names of abbreviations of brain regions are listed in **Supplementary Table S4**. (D) Correlation between memory response time and volume of the right scACC in FEP patients. (E) Region of interest representing the scACC overlaid in a medial sagittal section of a T1-weighted brain image in the Montreal Neurological Institute (MNI) space.

This result suggested that the relationship of these two areas, or both areas, or either area might have a particular signifiacence for this face processing measure. To address this exploratory question, we examined 46 rsFCs correlated (p-value < 0.05) with memory response time in FEP patients before multiple comparison correction. Interestingly, right scACC appeared much more prominently in this longer list (23 out of 46 rsFCs), whereas right precuneus only appeared in 2 rsFCs (**Fig. 2C**). To support this idea, when we examined a possible correlation bewteen scACC and memory response time, the volume of the right scACC was significantly correlated with this face processing measure in FEP patients (p-value = 0.02) (**Fig. 2D**). Together, our analysis pinned right scACC (**Fig. 2E**) as a specific brain region that is potentially involved in face processing.

## Discussion

In the present study, we addressed brain deficits associated with a dimension of social cognition in FEP patients by using the KDEF. By comparing FEP patients and HCs, we observed significant deficits in multiple face processing tasks that encompass *emotional processing* in FEP patients. We reproduced past reports in regard to impaired *emotional processing* in FEP patients (Barkl et al., 2014; Bozikas et al., 2019; Catalan et al., 2016; Dean et al., 2013), which may support the idea that the cohort and test battery used in the present study are compatible to other studies on social cognition deficits in psychosis. Importantly, we report correlations between face processing measures and the right scACC at both functional and structural levels in FEP patients.

There are multiple methods for defining sub-regions of the ACC. Multiple investigators have classically subdivided the ACC into the dorsal cognitive sub-region (BA32’ + BA24a’/b’/c’) and the rostral-ventral affective sub-region (BA32 + BA33 + BA24a/b + BA25) (Bush et al., 2000). Another method subdivided the ACC into four sub-regions: dorsal (BA24a’/b’/c’), rostral (rostral part of BA24a/b/c), subcallosal (caudal part of BA24a/b/c), and subgenual (BA25) (McCormick et al., 2006). Our segmentation based on the MRICloud included one specific segment equivalent to the subcallosal ACC sub-region defined by McCormick et al (McCormick et al., 2006). The present study underscored the unique involvement of this specific sub-region in face processing in FEP patients.

Three social cognitive networks (the mentalizing network, the action observation network, and the affective and value-based processing network) include the ACC in general (Apps et al., 2016). However, it is also well known that each sub-region of the ACC has a distinct function, connecting with different brain regions (Bush et al., 2000; McCormick et al., 2006). The main contribution of the present study is that we demonstrate the implication of scACC in face processing in FEP patients. Furthermore, we observed rsFCs between the scACC and precuneus. The precuneus has been involved in *theory of mind* (Baez et al., 2019) and our study now highlights this brain region in the context of *emotional processing*. As the precuneus is connected with ACC at the neurocircuitry level (Cavanna and Trimble, 2006), it may be reasonable that the rsFC between the scACC and this region showed significant correlation with *emotional processing*.

The scACC is reportedly involved in psychotic depression (Bijanki et al., 2014; McCormick et al., 2007). In addition to a significant reduction in scACC volume in patients with psychotic depression (Bijanki et al., 2014), low metabolic activity in the scACC was also shown in patients with psychotic depression (McCormick et al., 2007). Interestingly, the metabolic reduction seemed to be ameliorated after electroconvulsive therapy (McCormick et al., 2007). In regard to brain stimulation therapy for depression, BA25 has been underscored for deep brain stimulation (Mayberg et al., 2005; Rudebeck et al., 2019). Although BA25 is adjacent to the area we highlighted in the present study, the relationship between the scACC and BA25 in the context of depression remains to be elucidated. It is of interest how *emotional processing* assessed by KDEF may have implications in mood disorders possibly due to deficits of the scACC.

Human studies are useful in obtaining descriptive information that connects key behavioral constructs with specific brain regions/networks, but it is difficult to address their causal relationship. Animal models are expected to compliment this limitation. In studies with macaque, the ACC has been suggested to have a functional role in social cognition (Sallet et al., 2011; Sliwa and Freiwald, 2017). The ACC in macaque is anatomically and functionally comparable to human BA24a/b and BA24a’/b’ (Wittmann et al., 2018), which significantly overlap with the subcallosal part of BA24 (scACC) highlighted in the current study. The results of these macaque studies may further support the findings of the present human study. Reversely, the specific involvement of the scACC (caudal part of BA24a/b/c) may encourage finer experiments in macaque, in which the exact equivalent ACC sub-region is specifically studied for its causal involvement in *emotional processing* and social cognition.

The present study includes some points that we should cautiously consider. First, we may need to consider differences between the KDEF and the Penn Computerized Neurocognitive Battery/the Penn Emotion Recognition Task (Butler et al., 2009; Carter et al., 2009; Gur et al., 2017; Irani et al., 2012; Pinkham et al., 2018) that has been frequently used in psychotic study. For example, the KDEF includes four types of face expression (happy, angry, sad, and neutral), whereas the Penn battery includes five types of face expression (happy, angry, sad, fearful, and neutral). Furthermore, the KDEF includes pictures of only White individuals, whereas the Penn battery considers the balance of ethnicity. Thus, race was acknowledged as a possible confounding factor in our analysis, considering that the KDEF might cause other-race effect (Pinkham et al., 2008). However, when we compared performances between African American and other races, we did not observe significant differences in any domain. Our main conclusion may not be affected by the potential influence by race. Therefore, the concern regarding race may not affect the main conclusion of the present study. Second, response style of each participant may affect response time. Some participants may prioritize accuracy too much by sacrificing response time. Although response time is used as an outcome measure of face processing (Wilhelm et al., 2010), we need to be cautious in interpreting the data involving response time. Third, we regarded sex as a potential confounding factor in data analysis due to the small sample size of this cohort. However, in future studies with a larger sample size, it would be useful to consider biological effects of sex. Lastly, as described above, the correlation between face processing measures and multimodal MRI outcomes may not directly support their causal relationship. In order to resolve this issue, animal studies such as region-specific knockout models will be useful. Despite these points, we believe that the present study provides meaningful new evidence that a specific sub-region of the ACC (right subcallosal sub-region of BA24) is functionally and structurally important in FEP-associated pathological abnormalities in social cognition, in particular *emotional processing*.

## Funding

This work was supported by the National Institutes of Health (MH-094268, Silvio O. Conte Center, MH-092443, MH-105660, and MH-107730 to A.S., and S&R/RUSK, Stanley foundation. The Mitsubishi Tanabe Pharma Corporation supported a portion of the study participant recruitment cost.

## Notes

The authors wish to extend their gratitude to the participants in the current study. We appreciate Dr. Melissa A. Landek-Salgado for critical reading of this manuscript. We thank the study participant recruitment team led by Ms. Yukiko Lema. We also wish to thank Dr. Takanori Ogaru for their assistance in implementing the research.

## Notes

### Competing Interest Statement

The authors have declared no competing interest.

