## Supplementary materials for "Face processing of social cognition in patients with first episode psychosis: Its deficits and association with the right subcallosal anterior cingulate cortex"

Running title: Right subcallosal ACC and social cognition in FEP

Zui Narita,[1,6]|| Hironori Kuga,[1,6]|| Peeraya Piancharoen,[6] Andreia Faria,[2] Marina Mihaljevic,[1] Luisa Longo,[1] Semra Etyemez,[1,7] Ho Namkung,[1] Jennifer Coughlin,[1,2] Gerald Nestadt,[1] Frederik Nucifora,[1] Thomas Sedlak,[1] Rebecca Schaub,[1] Jeff Crawford,[1] David Schretlen,[1,2] Koko Ishizuka,[1] Jun Miyata,[8] Kun Yang,[1] and Akira Sawa[1,3,4,5,6]*

Departments of Psychiatry[1], Radiology and Radiological Sciences[2], Neuroscience[3], Biomedical Engineering[4], and Genetic Medicine[5],

Johns Hopkins University School of Medicine, Baltimore, Maryland.

Departments of Mental Health[6] and Epidemiology[7]

Johns Hopkins Bloomberg School of Public Health, Baltimore, Maryland.

Department of Psychiatry[8], Graduate School of Medicine, Kyoto University, Kyoto, Japan.

||These authors contributed equally to this work.

**Table S1. Two-group comparison of face processing measures between African American and non-African American subjects.**

A. healthy controls.

| Measures | p-value | q-value |
|---|---|---|
| Recognition accuracy | 0.44 | 0.44 |
| Recognition response time | 0.11 | 0.15 |
| Memory accuracy | 0.02 | 0.06 |
| Memory response time | 0.10 | 0.15 |

B. first episode psychosis patients.

| Measures | p-value | q-value |
|---|---|---|
| Recognition accuracy | 0.25 | 0.33 |
| Recognition response time | 0.11 | 0.21 |
| Memory accuracy | 0.04 | 0.14 |
| Memory response time | 0.76 | 0.76 |

**Table S2. Correlation between resting-state function connectivity (rsFC) and face processing measures in patients.**
The top 10 pairs were listed in each table. See abbreviations of symbols for brain regions in **Table S4**.

A. correlation between rsFCs and recognition accuracy.

| rsFC | correlation coefficient | p-value | q-value |
|---|---|---|---|
| ITG_R_LG_R | 0.40 | 2.05E-03 | 1.00 |
| SOG_R_Thalamus_R | -0.38 | 3.70E-03 | 1.00 |
| SOG_R_Thalamus_L | -0.37 | 4.76E-03 | 1.00 |
| IFG_opercularis_R_MTG_L_pole | -0.34 | 8.98E-03 | 1.00 |
| SOG_R_Cu_R | -0.34 | 8.99E-03 | 1.00 |
| ITG_R_LG_L | 0.34 | 9.36E-03 | 1.00 |
| ITG_R_SOG_R | 0.34 | 1.01E-02 | 1.00 |
| SFG_R_IFG_orbitalis_R | 0.33 | 1.08E-02 | 1.00 |
| MFOG_R_scACC_L | -0.33 | 1.11E-02 | 1.00 |
| ITG_R_Cu_R | 0.33 | 1.13E-02 | 1.00 |

B. correlation between rsFCs and recognition response time.

| rsFC | correlation coefficient | p-value | q-value |
|---|---|---|---|
| PoCG_L_Thalamus_L | 0.46 | 2.47E-04 | 0.36 |
| PrCG_L_Thalamus_L | 0.46 | 2.88E-04 | 0.36 |
| IFG_triangularis_L_scACC_R | 0.44 | 5.38E-04 | 0.36 |
| MFG_DPFC_R_scACC_R | 0.44 | 5.76E-04 | 0.36 |
| PoCG_L_Thalamus_R | 0.44 | 6.03E-04 | 0.36 |
| scACC_R_Caud_R | 0.42 | 1.09E-03 | 0.47 |
| PrCu_R_scACC_R | 0.42 | 1.09E-03 | 0.47 |
| IOG_R_Put_R | 0.40 | 1.64E-03 | 0.55 |
| scACC_R_Caud_L | 0.40 | 1.72E-03 | 0.55 |
| IFG_triangularis_R_scACC_R | 0.40 | 1.90E-03 | 0.55 |

(continue on the next page)

C. correlation between rsFCs and memory accuracy.

| rsFC | correlation coefficient | p-value | q-value |
|---|---|---|---|
| FuG_L_FuG_R | 0.50 | 7.26E-05 | 0.22 |
| MOG_R_IOG_R | 0.42 | 1.09E-03 | 1.00 |
| MOG_L_IOG_R | 0.41 | 1.49E-03 | 1.00 |
| FuG_R_MOG_L | 0.40 | 2.11E-03 | 1.00 |
| MFG_L_IFG_triangularis_L | -0.38 | 2.91E-03 | 1.00 |
| SFG_pole_R_PrCu_R | -0.35 | 6.64E-03 | 1.00 |
| FuG_R_SOG_R | 0.35 | 6.95E-03 | 1.00 |
| LFOG_R_MFOG_R | 0.35 | 7.02E-03 | 1.00 |
| FuG_R_MOG_R | 0.35 | 7.81E-03 | 1.00 |
| FuG_R_IOG_L | 0.34 | 9.98E-03 | 1.00 |

D. correlation between rsFCs and memory response time.

| rsFC | correlation coefficient | p-value | q-value |
|---|---|---|---|
| PrCu_R_scACC_R | 0.55 | 9.16E-06 | 0.03 |
| PrCu_L_scACC_R | 0.51 | 4.33E-05 | 0.07 |
| SPG_L_scACC_R | 0.48 | 1.21E-04 | 0.11 |
| MOG_R_scACC_R | 0.48 | 1.46E-04 | 0.11 |
| MOG_L_scACC_R | 0.46 | 2.71E-04 | 0.13 |
| SPG_R_scACC_R | 0.46 | 2.93E-04 | 0.13 |
| scACC_R_PCC_R | 0.46 | 3.08E-04 | 0.13 |
| MFG_R_IFG_triangularis_R | 0.45 | 4.54E-04 | 0.17 |
| MFG_L_IFG_triangularis_L | 0.42 | 1.14E-03 | 0.38 |
| MFG_R_IFG_orbitalis_R | 0.41 | 1.49E-03 | 0.43 |

**Table S3. Correlation between resting-state function connectivity (rsFC) and face processing measures in healthy controls.**
The top 10 pairs were listed in each table. See abbreviations of symbols for brain regions in **Table S4**.

A. correlation between rsFCs and recognition accuracy.

| rsFC | correlation coefficient | p-value | q-value |
|---|---|---|---|
| IFG_opercularis_L_Insula_R | -0.41 | 2.98E-04 | 0.82 |
| RG_L_dorsal_ACC_R | -0.39 | 5.44E-04 | 0.82 |
| LFOG_R_Thalamus_R | -0.38 | 8.54E-04 | 0.85 |
| IFG_orbitalis_L_IFG_triangularis_R | -0.37 | 1.31E-03 | 0.91 |
| IFG_orbitalis_L_Put_R | -0.36 | 1.86E-03 | 0.91 |
| LFOG_R_dorsal_ACC_R | -0.34 | 3.10E-03 | 0.91 |
| RG_L_SMG_L | -0.34 | 3.38E-03 | 0.91 |
| RG_L_Thalamus_R | -0.34 | 3.40E-03 | 0.91 |
| IFG_orbitalis_L_Thalamus_R | -0.34 | 3.45E-03 | 0.91 |
| RG_R_Put_L | -0.33 | 3.53E-03 | 0.91 |

B. correlation between rsFCs and recognition response time.

| rsFC | correlation coefficient | p-value | q-value |
|---|---|---|---|
| SOG_R_Cu_R | -0.31 | 6.95E-03 | 1.00 |
| SFG_L_CerebellumGM_L | 0.31 | 7.65E-03 | 1.00 |
| IFG_opercularis_L_MTG_L | -0.31 | 8.14E-03 | 1.00 |
| dorsal_ACC_R_CerebellumGM_L | 0.30 | 1.04E-02 | 1.00 |
| PrCu_R_CerebellumGM_R | 0.30 | 1.04E-02 | 1.00 |
| PCC_R_CerebellumGM_L | 0.29 | 1.08E-02 | 1.00 |
| PrCu_R_CerebellumGM_L | 0.29 | 1.37E-02 | 1.00 |
| MTG_L_pole_subgenual_ACC_R | 0.28 | 1.62E-02 | 1.00 |
| dorsal_ACC_R_CerebellumGM_R | 0.28 | 1.70E-02 | 1.00 |
| ITG_L_Cu_R | 0.28 | 1.71E-02 | 1.00 |

(continue on the next page)

C. correlation between rsFCs and memory accuracy.

| rsFC | correlation coefficient | p-value | q-value |
|---|---|---|---|
| SPG_R_subgenual_ACC_L | 0.44 | 7.58E-05 | 0.23 |
| FuG_L_SOG_L | 0.36 | 1.60E-03 | 0.98 |
| STG_R_Cu_R | 0.35 | 2.32E-03 | 0.98 |
| FuG_L_subgenual_ACC_L | 0.35 | 2.43E-03 | 0.98 |
| RG_L_SPG_R | 0.34 | 2.89E-03 | 0.98 |
| SPG_L_subgenual_ACC_L | 0.33 | 3.84E-03 | 0.98 |
| RG_R_SPG_R | 0.33 | 3.85E-03 | 0.98 |
| scACC_R_Thalamus_L | -0.31 | 6.47E-03 | 0.98 |
| PoCG_R_STG_L_pole | 0.31 | 7.95E-03 | 0.98 |
| MFG_L_IFG_triangularis_L | -0.30 | 8.67E-03 | 0.98 |

D. correlation between rsFCs and memory response time.

| rsFC | correlation coefficient | p-value | q-value |
|---|---|---|---|
| IFG_triangularis_L_SMG_L | -0.34 | 3.05E-03 | 1.00 |
| MFG_DPFC_R_IFG_triangularis_R | -0.31 | 7.98E-03 | 1.00 |
| Cu_R_LG_L | -0.30 | 9.16E-03 | 1.00 |
| PrCG_R_STG_R | -0.30 | 9.83E-03 | 1.00 |
| FuG_L_scACC_L | -0.29 | 1.13E-02 | 1.00 |
| MFG_DPFC_L_IFG_orbitalis_L | -0.29 | 1.16E-02 | 1.00 |
| IFG_opercularis_L_IFG_triangularis_L | -0.29 | 1.34E-02 | 1.00 |
| SMG_R_scACC_L | -0.28 | 1.56E-02 | 1.00 |
| RG_L_ITG_L | -0.28 | 1.56E-02 | 1.00 |
| MFG_L_MTG_L | -0.28 | 1.58E-02 | 1.00 |

**Table S4. Abbreviations of symbols for brain regions.**

| abbreviation | region |
|---|---|
| _L | Left Hemisphere |
| _R | Right Hemisphere |
| AG | Angular gyrus |
| Caud | Caudate nucleus |
| CerebellumGM | Cerebellum gray matter |
| Cu | Cuneus |
| dorsal_ACC | Dorsal anterior cingulate gyrus |
| FuG | Fusiform gyrus |
| IFG_opercularis | Inferior frontal gyrus pars opercularis |
| IFG_orbitalis | Inferior frontal gyrus pars orbitralis |
| IFG_triangularis | Inferior frontal gyrus pars triangularis |
| Insula | Insula |
| IOG | Inferior occipital gyrus |
| ITG | Inferior temporal gyrus |
| LFOG | Lateral fronto-orbital gyrus |
| LG | Lingual gyrus |
| MFG | Middle frontal gyrus |
| MFG_DPFC | Middle frontal gyrus |
| MFOG | Middle fronto-orbital gyrus |
| MOG | Middle occipital gyrus |
| MTG | Middle temporal gyrus |
| PCC | Posterior cingulate gyrus |
| PoCG | Postcentral gyrus |
| PrCG | Precentral gyrus |
| PrCu | Pre-cuneus |
| Put | Putamen |
| RG | Gyrus rectus |
| SFG | Superior frontal gyrus |
| SFG_PFC | Superior frontal gyrus |
| SMG | Supramarginal gyrus |
| SOG | Superior occipital gyrus |
| SPG | Superior parietal gyrus |
| STG | Superior temporal gyrus |
| scACC | Subcallosal anterior cingulate gyrus |
| subgenual_ACC | Subgenual anterior cingulate gyrus |
| Thalamus | Thalamus |
